# Spatial distribution and factors associated with low birth weight in Ethiopia using data from Ethiopian demographic and health survey 2016: spatial and multilevel analysis

**DOI:** 10.1101/2020.06.04.134007

**Authors:** Alemneh Mekuriaw Liyew, Malede Mequanent Sisay, Achenef Asmamaw Muche

## Abstract

**Background:** Low birth weight (LBW) was a leading cause of neonatal mortality. It showed an increasing trend in Sub-Saharan Africa for the last one and half decade. Moreover, it was a public health problem in Ethiopia. Even though different studies were conducted to identify its predictors, contextual factors were insufficiently addressed in Ethiopia. There was also limited evidence on the spatial distribution of low birth weight. Therefore, this study aimed to explore spatial distribution and factors associated with low birth weight in Ethiopia.

**Method:** Secondary data analysis was conducted using the 2016 EDHS data. A total of 1502 (weighted sample) mothers whose neonates were weighed at birth five years preceding the survey were included. GIS 10.1, SaTscan, stata, and Excel were used for data cleaning and analysis. A multi-level mixed-effects logistic regression model was fitted to identify factors associated with low birth weight. Finally, hotspot areas from GIS results, log-likelihood ratio (LLR) and relative risk with p-value of spatial scan statistics, AOR with 95% CI and random effects for mixed-effects logistic regression model were reported.

**Results:** Low birth weight was spatially clustered in Ethiopia. Primary (LLR=11.57; P=0.002) clusters were detected in the Amhara region. Whereas secondary (LLR=11.4; P=0.003;LLR=10.14,P=0.0075) clusters were identified at Southwest Oromia, north Oromia, south Afar, and Southeast Amhara regions. Being severely anemic (AOR=1.47;95%CI1.04,2.01), having no education (AOR=1.82;95%CI1.12,2.96), Prematurity (AOR=5.91;95%CI3.21,10.10) female neonate (AOR=1.38;95%CI1.04,1.84)were significantly associated with LBW

**Conclusion:** LBW was spatially clustered in Ethiopia with high-risk areas in Amhara,Oromia, and Afar regions and it was affected by socio demographic factors. Therefore, focusing the policy intervention in those geogrsphically low birth weight risk areas and improving maternal education and nutrtion could be vital to reduce the low birth weight disparity in Ethiopia.

## Background

World health organization defined low birth weight (LBW) as weight at birth less than 2,500 grams (1). It was strongly linked to neonatal mortality. Globally neonatal deaths accounted for 46% of all under-five deaths of which 38% of newborn deaths occurred in sub-Saharan Africa. Ethiopia was the one among five countries those account for about half of all global neonatal deaths (2). The 2016 EDHS report showed that the neonatal mortality rate was 29/1000 live births and Ethiopia fails to achieve the millennium development goal target for neonatal mortality (2, 3)..

Low birth weight directly or indirectly accounted for about 60-80% of neonatal mortality in Asia (4) and LBW babies were highly Vulnerable to death than heavier babies (5). Besides, as birth weight decreases the mortality increases ranging up to 100 fold variation across the birth weight spectrum. In Ethiopia, Low birth weight accounted for 3.63% of total deaths (6).

The consequence of LBW was not limited to neonatal and infant mortality but it also results in physical and developmental health problems in subsequent childhood and adulthood life. It leads to poor childhood growth and a higher incidence of adulthood chronic diseases like type 2 diabetes, hypertension and cardiovascular disease (7). This showed that LBW was a basement for the majority of adulthood chronic diseases. It has also long term consequences like poor cognitive function, academic underachievement and impaired behavior(8, 9). Moreover, LBW was a summary measure of multifaceted public health problems such as maternal malnutrition, ill-health and poor pregnancy-related health service utilization (7, 10).

Globally, more than 20 million infants were born being LBW. Among these nearly half (48%) of LBW, births occurred in southern Asia. In sub-Saharan Africa, the number of LBW live births was estimated to have increased from 4·4 million in 2000 to 5·0 million in 2015 (11). Furthermore, LBW was continuing as a public health challenge in this region (12). It was worth noting that these rates were high, even though the data on low birth weight remain limited as many deliveries occur at home or small health clinics and were not reported in official figures, which may result in an underestimation of the prevalence. Therefore World health organization incorporated as a third target to achieve a 30% reduction in low birth weight incidence by 2025 (13).

Even though data on low birth weight was limited due to low institutional delivery in Ethiopia, the prevalence of low birth weight was increased by 5% from 2000 (14) to 2016 (15). Furthermore, the evidence from systematic review and meta-analysis showed that the pooled prevalence of low birth weight to be 17.3% (16).

Spatial studies conducted in different middle and upper-income countries (17) showed that LBW was spatially clustered in different regions of the respective countries. In Ethiopia, the prevalence of LBW was different across different geopolitical regions (18–20). This showed the variation in the prevalence of LBW across different administrative regions provides an insight to identify risk (hotspot) areas by using spatial technology.

A research conducted in different countries on determinants of LBW showed that various socio-demographic, socioeconomic, pregnancy and maternal health service-related and community-related factors to be predictors of low birth weight (18, 21, 22). In Ethiopia, prior studies have been done to identify sociodemographic, pregnancy, and maternal health service-related factors (20–23). Even though the low birth weight was affected by factors operating at both individual and community level none of the studies have tried to look at the factors that affect LBW at community and individual levels simultaneously. Furthermore, there was limited evidence on the spatial distribution of low birth weight in Ethiopia.

Therefore, this study was aimed at identifying both individual and community-level factors associated with low birth weight simultaneously by applying a multi-level modeling approach. Besides, it tried to identify high risk (hot spot areas)of low birth weight by applying spatial analysis to provide evidence for socio-demographic to have a better understanding.

## Methods

### Data source

Secondary data analysis was employed to identify spatial distribution and factors associated with low birth weight. An authorization letter for the use of this data was obtained from Measure DHS and the dataset was downloaded from measure DHS website www.measuredhs.com.The survey covered all the nine regions and two administrative cities in Ethiopia (Fig.1).

**Fig. 1.**
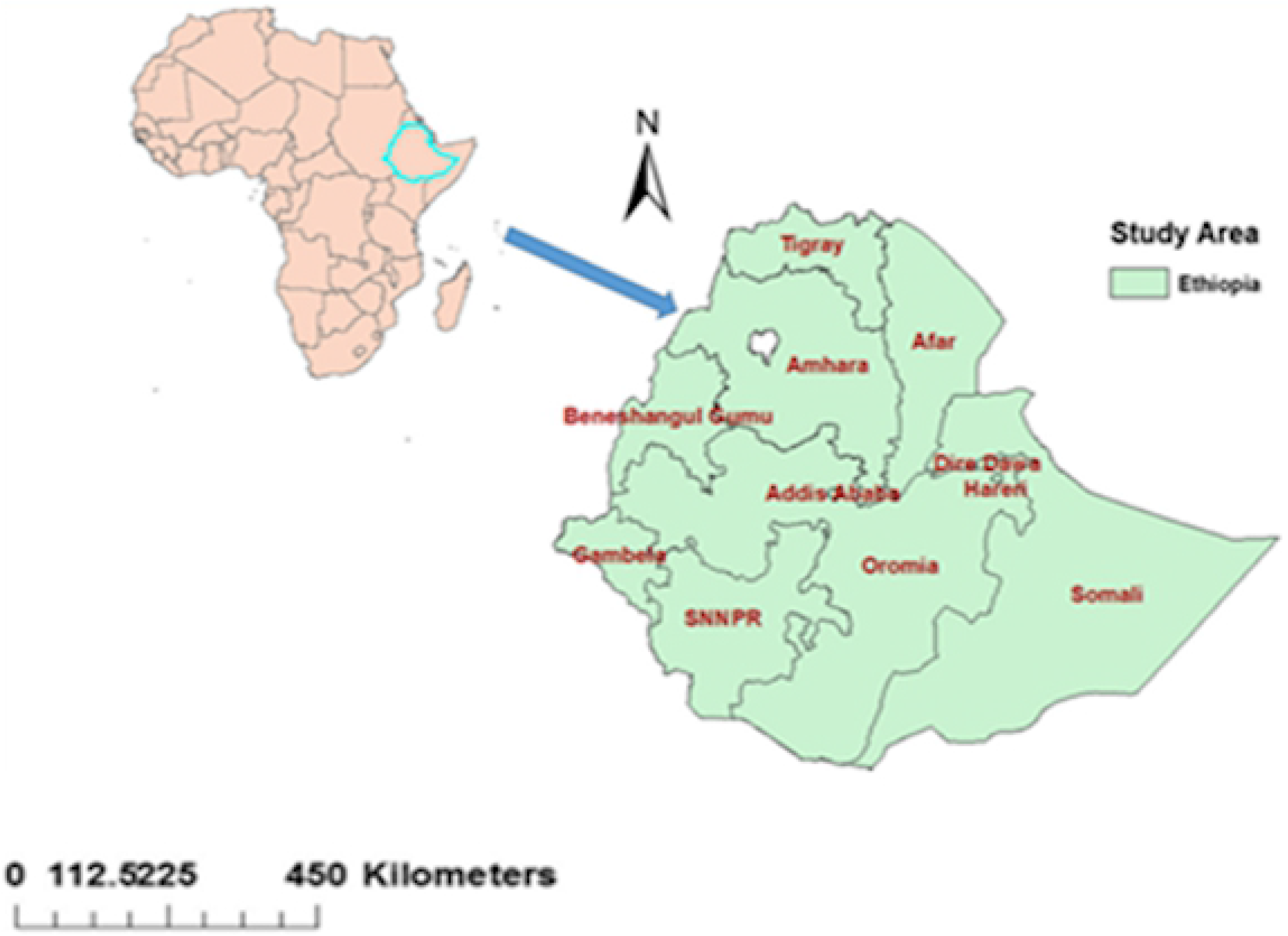
Map of Ethiopia with nine regions and two administrative cities as labeled on the map.

The participants were selected using a stratified two-stage cluster sampling technique. The survey collected information from a nationally representative sample of 16583 eligible women within 645 enumeration areas. The full method applied to the data collection procedure for EDHS 2016 was published elsewhere (15). For this study, to adjust for over or under-sampling which might occur due to the sampling nature of Demographic and health survey we applied sampling weight. This was considered to produce a nationally representative sample since those regions with a larger population could be under-sampled whereas those regions with a small population could be oversampled whenever sampling weight was not applied. Therefore, the final weighted sample size for our analysis was 1502 neonates born five years preceding the survey nested within 542 communities (clusters) for multilevel analysis. Since clusters with no recorded geographic coordinate were excluded, the spatial analysis was based on 441 clusters. The weighted proportion of LBW per cluster was computed for further spatial analysis.

### Study Variables

#### Dependent variable

The main outcome variable of this study was birth weight. Data on the birth weight of children were collected from mothers who gave birth within 5 years before the survey either by accessing birth weight through record review or by the mother’s report by recalling the measured weight of the child at birth. The births without recorded birth weight were excluded from the study. Finally, it was categorized as birth weight ≥2.5kg or <2.5 kg.

#### Independent variables

The determinants of low birth weight were extracted after reviewing literature at a global level. Maternal Age, maternal Educational status, sex of infant, wealth index, media exposure, number of ANC visit, gestational age, Maternal anemia, Maternal BMI, Iron supplementation, maternal height, birth order, birth interval, and cesarean delivery were individual-level predictors. Whereas, community poverty, community media exposure, community women education, region, and place of residence were community-level variables.

The aggregate community level explanatory variables were constructed by aggregating individual-level characteristics at the community (cluster) level. They were dichotomized as high or low based on the distribution of the proportion values computed for each community after checking the distribution by using the histogram. If the aggregate variable was normally distributed mean value and if not, normally distributed median value was used as a cut-off point for the categorization. Community poverty level was categorized as high if the proportion of women from the two lowest wealth quintiles in a given community was 1–100 % and low if the proportion was 0 %. Community media exposure was categorized as low if the proportion of women exposed to media in the community was 0–83.33 % and categorized as high if the proportion was 83.33–100 %, Community Women education was categorized as low if the proportion of women with no formal education in the community was 14.83%-100% and categorized as high if the proportion was 0–14.83 %.

### Spatial analysis

#### Spatial autocorrelation analysis

The Global Moran’s I statistic test was used to measure whether the low birth weight patterns were randomly distributed, dispersed, or clustered in Ethiopia. The calculated Moran’s I values close to −1 indicate disease dispersed, whereas I close to +1 indicate disease clustered and if the I value zero the disease is distributed randomly A statistically significant Moran’s I (p < 0.05) leads to rejection of the null hypothesis(random distribution of the diseases in the study area) and implies the presence of spatial autocorrelation (23, 24).

#### Spatial scan statistical analysis

Spatial Scan statistical method is commonly recommended that it is better than others in detecting local clusters and has higher power as compared to available spatial statistical methods (25).

The presence of statistically significant spatial hotspots/clusters of low birth weight) was tested by using spatial scan statistical analysis. It uses a scanning window that moves across the study area (26, 27). Low birth weight newborns were taken as cases and those were not born being LBW were taken as controls to fit the Bernoulli model. The number of cases in each location had Bernoulli distribution and the model requires data with or without a disease.

The default maximum spatial cluster size of <50% of the population was used, as an upper limit, which allowed both small and large clusters to be detected and ignored clusters that contained more than the maximum limit. For each potential cluster, a likelihood ratio test statistic was used to determine if the number of observed LBW cases within the potential cluster was significantly higher than expected or not. The primary and secondary clusters were identified and assigned p-values and ranked based on their likelihood ratio test, based on 999 Monte Carlo replications (28, 29).

#### Spatial interpolation

It is very expensive and laborious to collect reliable data in all areas of the country to know the burden of certain events. Therefore, part of a certain area can be predicted by using observed data using a method called interpolation. Spatial interpolation technique used to predict low birth weight on the un-sampled areas in the country based on sampled enumeration areas. There are various deterministic and geostatistical interpolation methods. Among all of the methods, ordinary Kriging and empirical Bayesian Kriging are considered the best method since it incorporates the spatial autocorrelation and it statistically optimizes the weight. For this study, the ordinary Kriging method was used to estimate the burden of low birth weight in unsampled areas.

#### Multi-level analysis

The outcome variable was dichotomized as birth weight <2.5Kg and ≥2.5Kg. Sampling weight was applied as part of a complex survey design using primary sampling unit, strata and women’s individual weight (V005).

Because of the hierarchical nature of data and the dichotomous outcome variable, the multi-level mixed-effects logistic regression model was used. The individual and community level variables associated with LBW were checked independently in the bivariable multilevel logistic regression model and Variables which were statistically significant at p-value 0.2 in the bivariable multilevel mixed-effects logistic regression analysis were considered for the individual and community level model adjustments.

#### Model building

Four models were fitted. The first was the null model containing no exposure variables which was used to check variation in community and provide evidence to assess random effects at the community level. The second model was the multivariable model adjustment for individual-level variables and model three was adjusted for community-level factors. In the fourth model, possible candidate variables from both individual and community-level variables were fitted with the outcome variable.

#### Parameter estimation method

The fixed effects (measure of association) were used to estimate the association between the likelihood of low birth weight and explanatory variables at both community and individual level and were expressed as odds ratio with 95% confidence interval. Regarding the measures of variation (random-effects) intracluster correlation coefficient (ICC), Proportional Change in Community Variance (PCV), and median odds ratio (MOR) were used.

## Results

In this study total of 1502(weighted) women whose neonate was weighed at birth were included. Of all participants, 28.84% had no formal education and the Mean age of the mothers was 28.5[SD=±6.08] years. About 67% of participants were exposed to media. Regarding anemia and iron supplementation, 14.31% of the participants were severely anemic and 4 3.21% didn’t receive iron during pregnancy(see Table1).

**Table 1:**
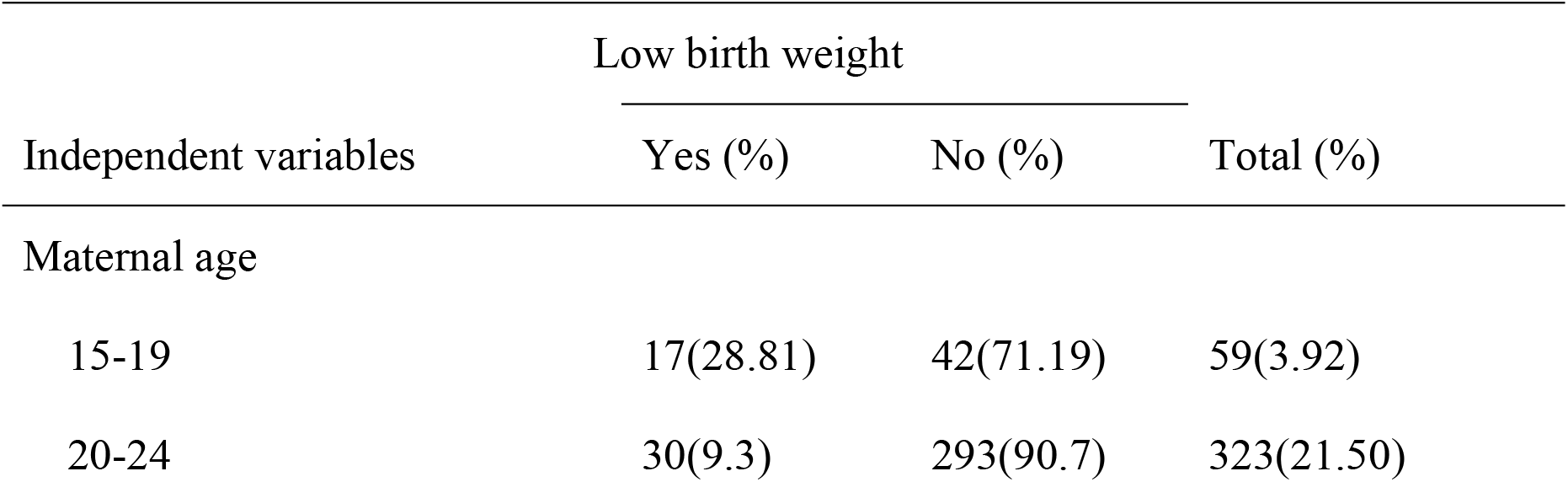
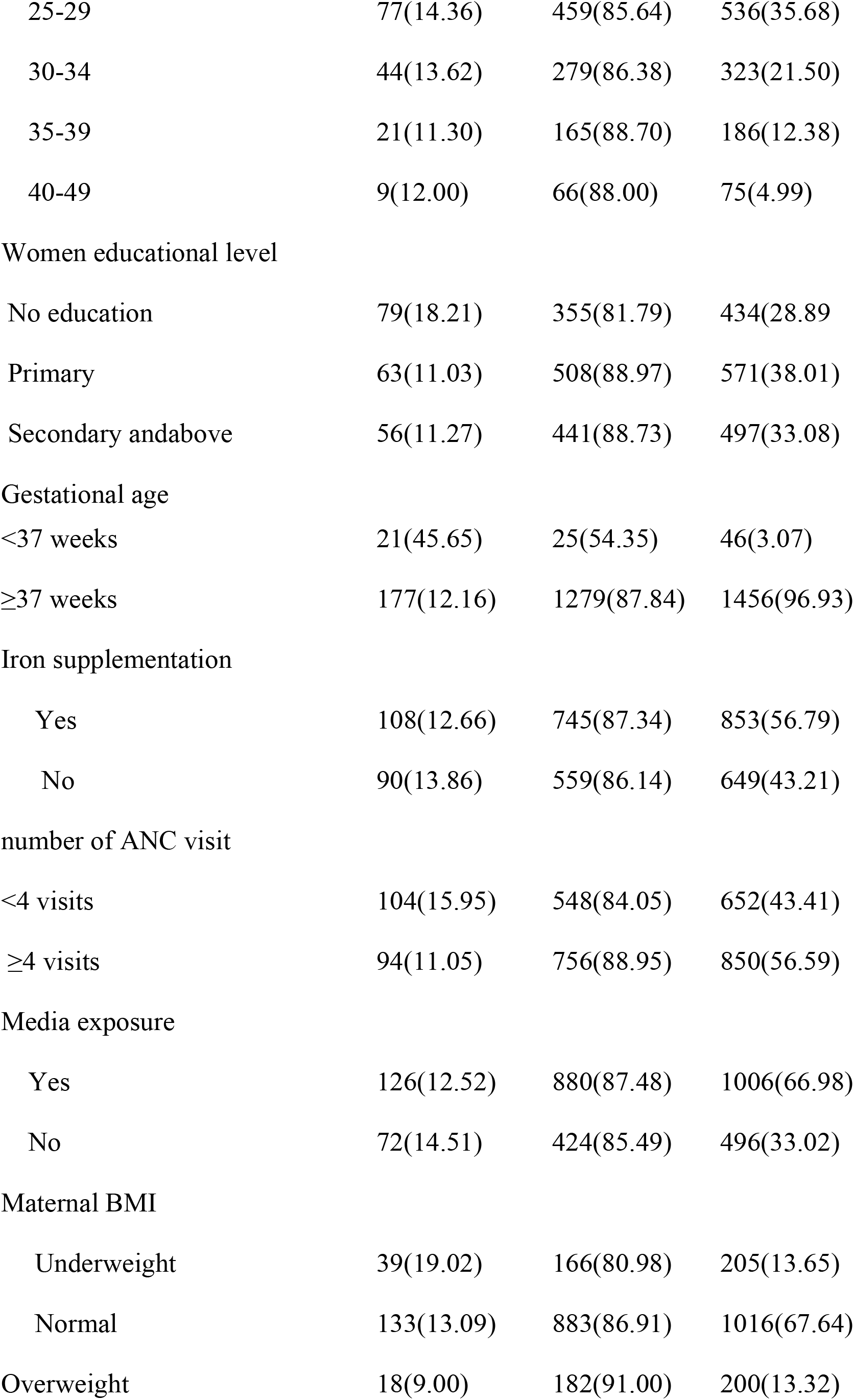
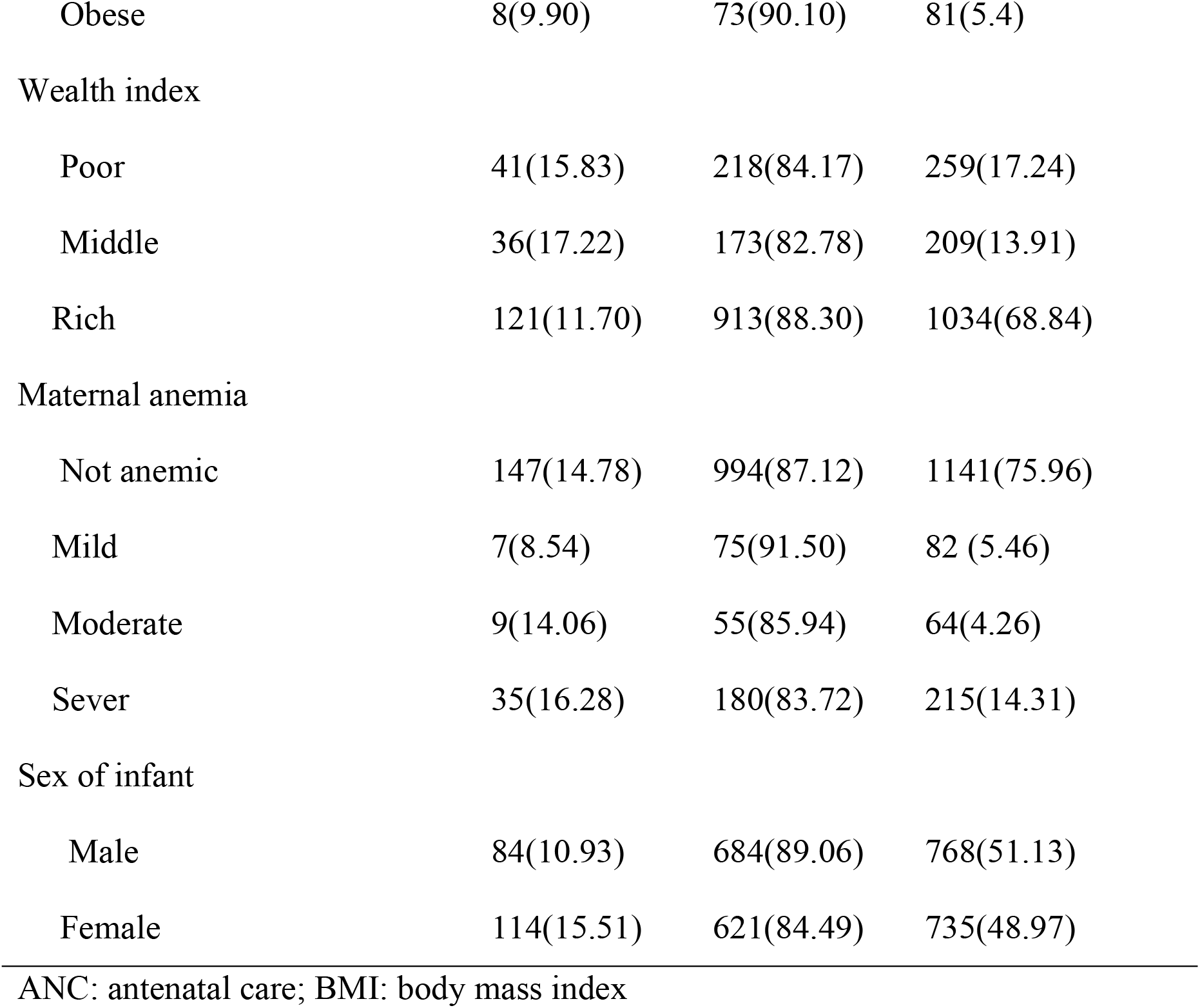
Individual level characteristics of women who give live birth in Ethiopia, EDHS 2016 perspective (N=1502 weighted).

| Independent variables | Low birth weight |  | Total (%) |
| --- | --- | --- | --- |
|  | Yes (%) | No (%) |  |
| Maternal age |  |  |  |
| 15-19 | 17(28.81) | 42(71.19) | 59(3.92) |
| 20-24 | 30(9.3) | 293(90.7) | 323(21.50) |
| 25-29 | 77(14.36) | 459(85.64) | 536(35.68) |
| 30-34 | 44(13.62) | 279(86.38) | 323(21.50) |
| 35-39 | 21(11.30) | 165(88.70) | 186(12.38) |
| 40-49 | 9(12.00) | 66(88.00) | 75(4.99) |
| Women educational level |  |  |  |
| No education | 79(18.21) | 355(81.79) | 434(28.89) |
| Primary | 63(11.03) | 508(88.97) | 571(38.01) |
| Secondary and above | 56(11.27) | 441(88.73) | 497(33.08) |
| Gestational age |  |  |  |
| <37 weeks | 21(45.65) | 25(54.35) | 46(3.07) |
| ≥37 weeks | 177(12.16) | 1279(87.84) | 1456(96.93) |
| Iron supplementation |  |  |  |
| Yes | 108(12.66) | 745(87.34) | 853(56.79) |
| No | 90(13.86) | 559(86.14) | 649(43.21) |
| number of ANC visit |  |  |  |
| <4 visits | 104(15.95) | 548(84.05) | 652(43.41) |
| ≥4 visits | 94(11.05) | 756(88.95) | 850(56.59) |
| Media exposure |  |  |  |
| Yes | 126(12.52) | 880(87.48) | 1006(66.98) |
| No | 72(14.51) | 424(85.49) | 496(33.02) |
| Maternal BMI |  |  |  |
| Underweight | 39(19.02) | 166(80.98) | 205(13.65) |
| Normal | 133(13.09) | 883(86.91) | 1016(67.64) |
| Overweight | 18(9.00) | 182(91.00) | 200(13.32) |
| Obese | 8(9.90) | 73(90.10) | 81(5.4) |
| Wealth index |  |  |  |
| Poor | 41(15.83) | 218(84.17) | 259(17.24) |
| Middle | 36(17.22) | 173(82.78) | 209(13.91) |
| Rich | 121(11.70) | 913(88.30) | 1034(68.84) |
| Maternal anemia |  |  |  |
| Not anemic | 147(14.78) | 994(87.12) | 1141(75.96) |
| Mild | 7(8.54) | 75(91.50) | 82 (5.46) |
| Moderate | 9(14.06) | 55(85.94) | 64(4.26) |
| Sever | 35(16.28) | 180(83.72) | 215(14.31) |
| Sex of infant |  |  |  |
| Male | 84(10.93) | 684(89.06) | 768(51.13) |
| Female | 114(15.51) | 621(84.49) | 735(48.97) |

ANC: antenatal care; BMI: body mass index

### Community level characteristics of participants

A total of 452 communities (clusters) were included in this study. About two-thirds (67.44%) of mothers were from a community with a low poverty level. About half (51.33%) of participants were from rural communities (see table2).

**Table 2:**
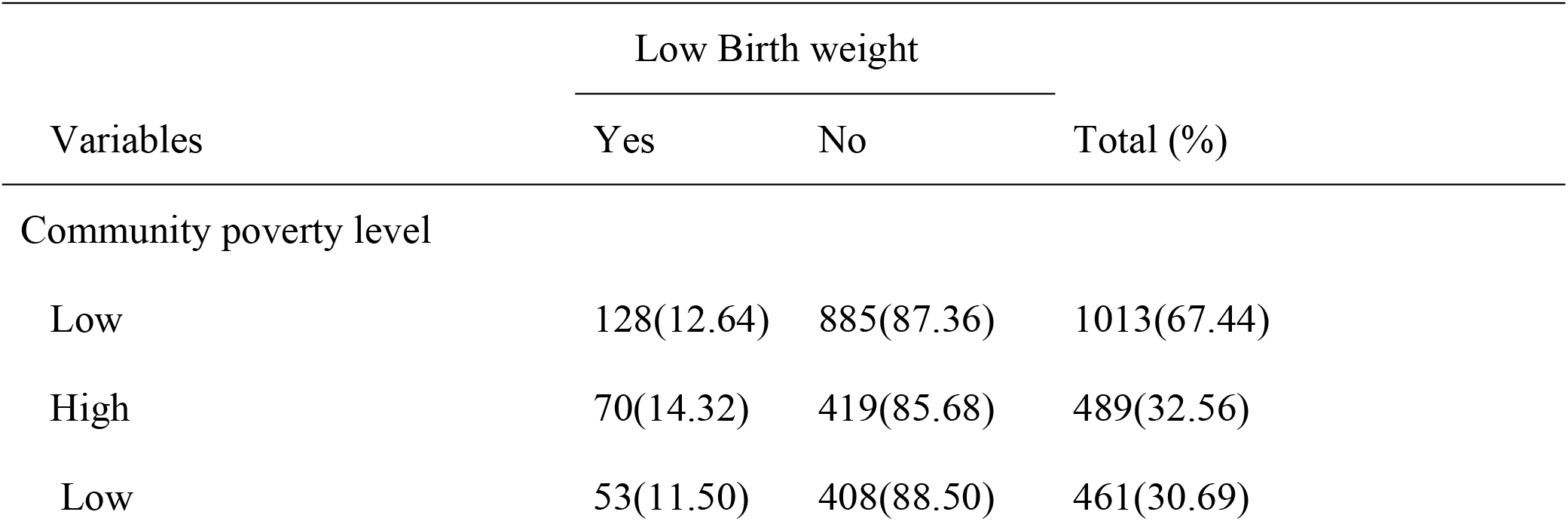
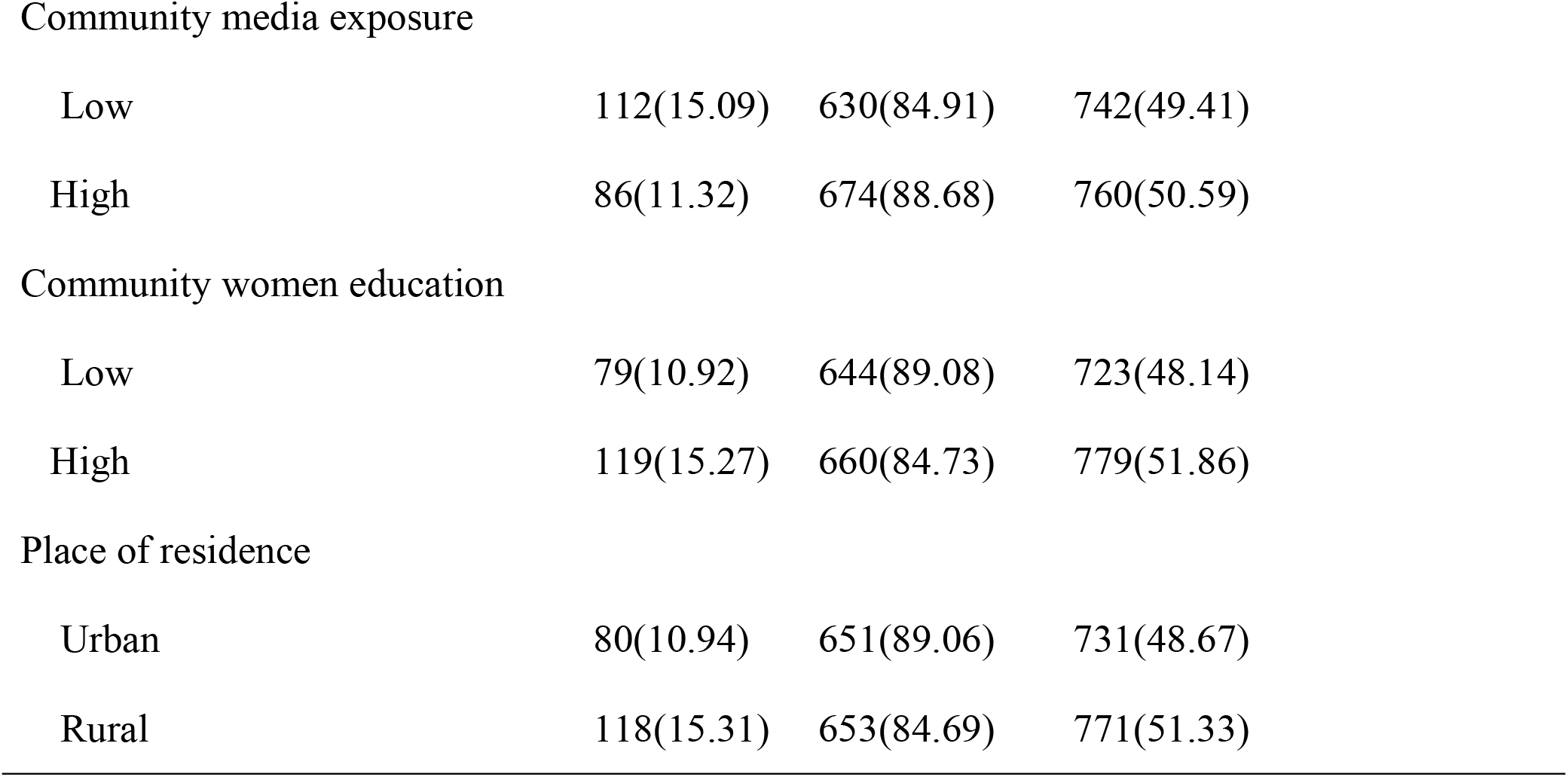
Community level characteristics of women EDHS 2016 (N=1502).

| Variables | Low Birth weight |  | Total (%) |
| --- | --- | --- | --- |
|  | Yes | No |  |
| Community poverty level |  |  |  |
| Low | 128(12.64) | 885(87.36) | 1013(67.44) |
| High | 70(14.32) | 419(85.68) | 489(32.56) |
| Low | 53(11.50) | 408(88.50) | 461(30.69) |
| Community media exposure |  |  |  |
| Low | 112(15.09) | 630(84.91) | 742(49.41) |
| High | 86(11.32) | 674(88.68) | 760(50.59) |
| Community women education |  |  |  |
| Low | 79(10.92) | 644(89.08) | 723(48.14) |
| High | 119(15.27) | 660(84.73) | 779(51.86) |
| Place of residence |  |  |  |
| Urban | 80(10.94) | 651(89.06) | 731(48.67) |
| Rural | 118(15.31) | 653(84.69) | 771(51.33) |

### Spatial distribution of low birth weight in Ethiopia

The spatial variation of proportion of low birth weight was maped. Thus, the high prevalence of low birth weight was observed in Afar, North-west Amhara, Northeast SNNP, central part of Oromia region and Somali regional states of Ethiopia. (Fig 2).

**Fig. 2.**
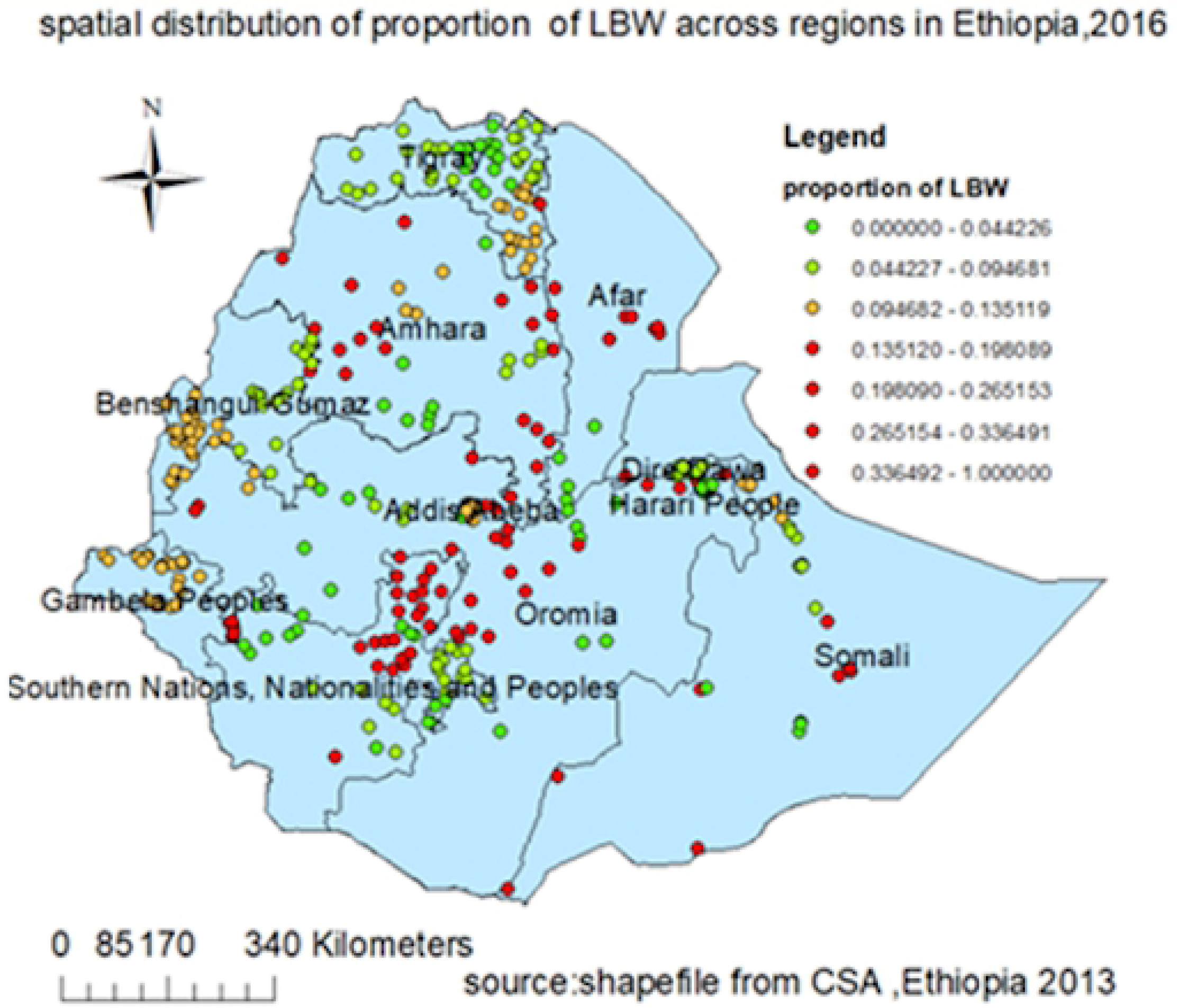
Spatial distribution of low birth weight in Ethiopia 2016. In this figure each spot on the map represents single enumeration area. Red dots show enumeration areas with higher prevalence of LBW while green ones show lower prevalence of LBW.

Fig 2: Spatial distribution of low birth weight in Ethiopia.

### Spatial autocorrelation of low birth weight in Ethiopia

This study identified that the spatial distribution of low birth weight was found to be clustered in Ethiopia with Global Moran’s I =0.56 and p= 0.001 (Fig 3). The clustered patterns (on the right sides) show high rates of low birth weight occurred over the study area. The Z-score of 45.57 indicated that there is less than 1% likelihood that this clustered pattern could be the result of random chance.

**Fig.3.**
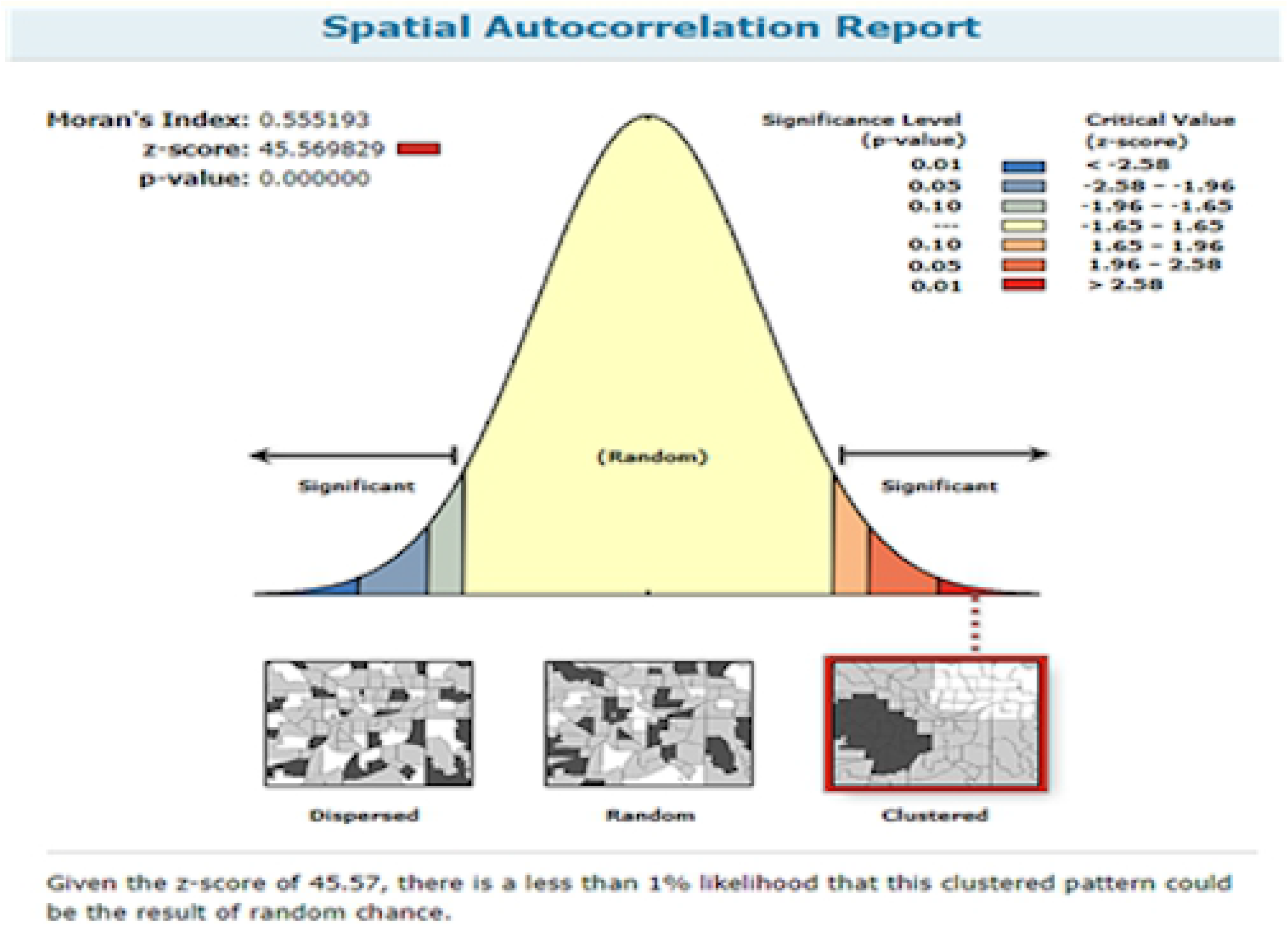
Spatial autocorrelation analysis of LBW in Ethiopia 2016. The clustered pattern (on the right sides) shows high rates of LBW occurred over the study area. Auto-generated interpretations available underneath show that the likelihood that clustered pattern could be due to random chance is less than 1 %. The bright red and blue color (to the end tails) indicates increased significance level.

Fig 3: Spatial autocorrelation of low birth weight in Ethiopia, EDHS 2016.

### Spatial scan statistics of low birth weight in Ethiopia

Spatial scan statistics identified 40 significant clusters of which 15 were primary clusters and 25 were secondary clusters. The primary clusters’ spatial window was located in the North West Amhara and Northeast part of BenishangulGumuz, which was centered at 11.57418N, 36.498123 E with 122.56km radius, and Log-Likelihood ratio (LLR) of 11.82, at p < 0.01. It showed that women within the spatial window had 2.66 times higher risk of LBW as compared to women outside the window.

Two other significant spatial windows were located in southwest Oromia and at the border of southeast Amhara, south Afar and northern part of Oromia regions. The one located in southwest Oromia region was centered at 7.192884N,39.02565E with 30.83 radius, LLR of 11.43 and p-value 0. 003. Women within this scanning window had 4.82 times higher risk of LBW than those outside the scanning window. The third scanning window which was located at the border of three regions (Amhara,Afar and Oromia) was centered at 10.143320N,39.718498E with 158.43km radius and log-likelihood ratio (LLR) of 10.37 at p-value 0. 0075. It showed that women within the window had 2.52 times higher risk of LBW as compared to those outside the window. (Fig 4, Table 3).

**Table 3.** significant spatial clusters of low birth weight in Ethiopia EDHS 2016.

| clusters | Enumeration areas detected | population | cases | LLR | RR | Coordinate/radius | p-value |
| --- | --- | --- | --- | --- | --- | --- | --- |
| 1* | 259,602,541,386,548,361,515,498,516,109,292,533,73,167,52 | 94 | 30 | 11.82 | 2.66 | 11.574184N,36.498123E/122.56km | 0.0021 |
| 2** | 26,589 | 18 | 11 | 11.41 | 4.82 | 7.192884 N, 39.025650 E) / 30.83 km | 0.003 |
| 3** | 637, 310, 295, 484, 624, 135, 617, 230, 18, 423, 121, 616, 49, 345, 254, 611, 71, 40, 303, 90, 287, 402, 560 | 95 | 29 | 10.37 | 2.52 | 10.143320 N, 39.718498 E) / 158.43 km | 0.0075 |

\*=primary clusters;\*\*=secondary clusters

**Fig.4.**
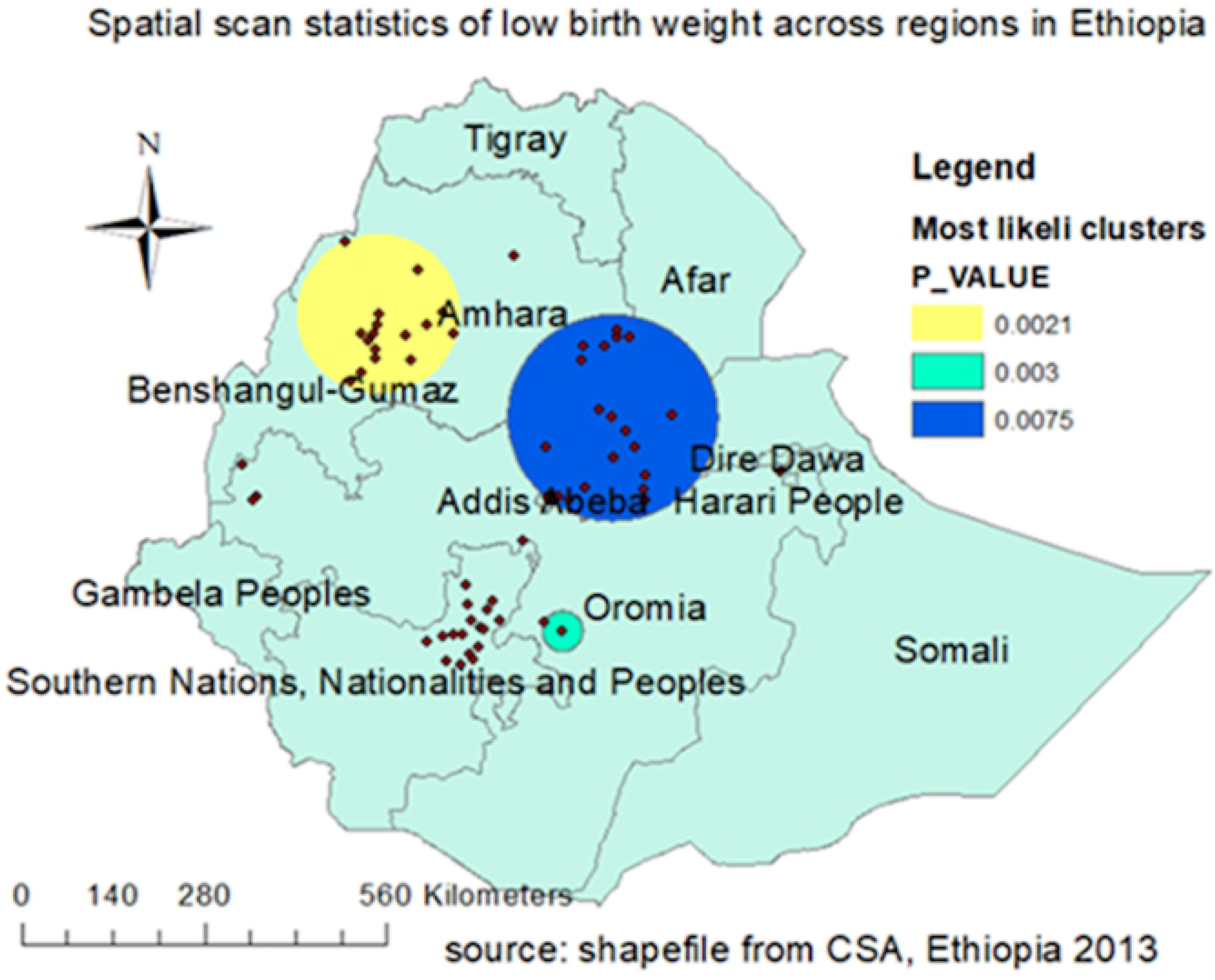
Most likely and secondary clusters of LBW in Ethiopia, EDHS 2016. TI1e bright blue colors (rings) indicate the most statistically significant spatial windows which contain primary clusters of LBW. Women within the spatial window had higher risk of bearing LBW baby than those outside the window.

Fig. 6: The spatial scaning statistcs of LBW in Ethiopia, 2016.

### Spatial interpolation of LBW in Ethiopia in 2016

In ordinary kriging spatial interpolation zone1, zone4 and zone 5 in Afar region; Waghimra, Semen Gonder, and Awi in Amhara region; mirabwelega and mirabArsi in Oromia region; Gurage, silti and debubomo in SNNP region were areas at high-risk of low birth weight (Fig. 5).

**Fig.5.**
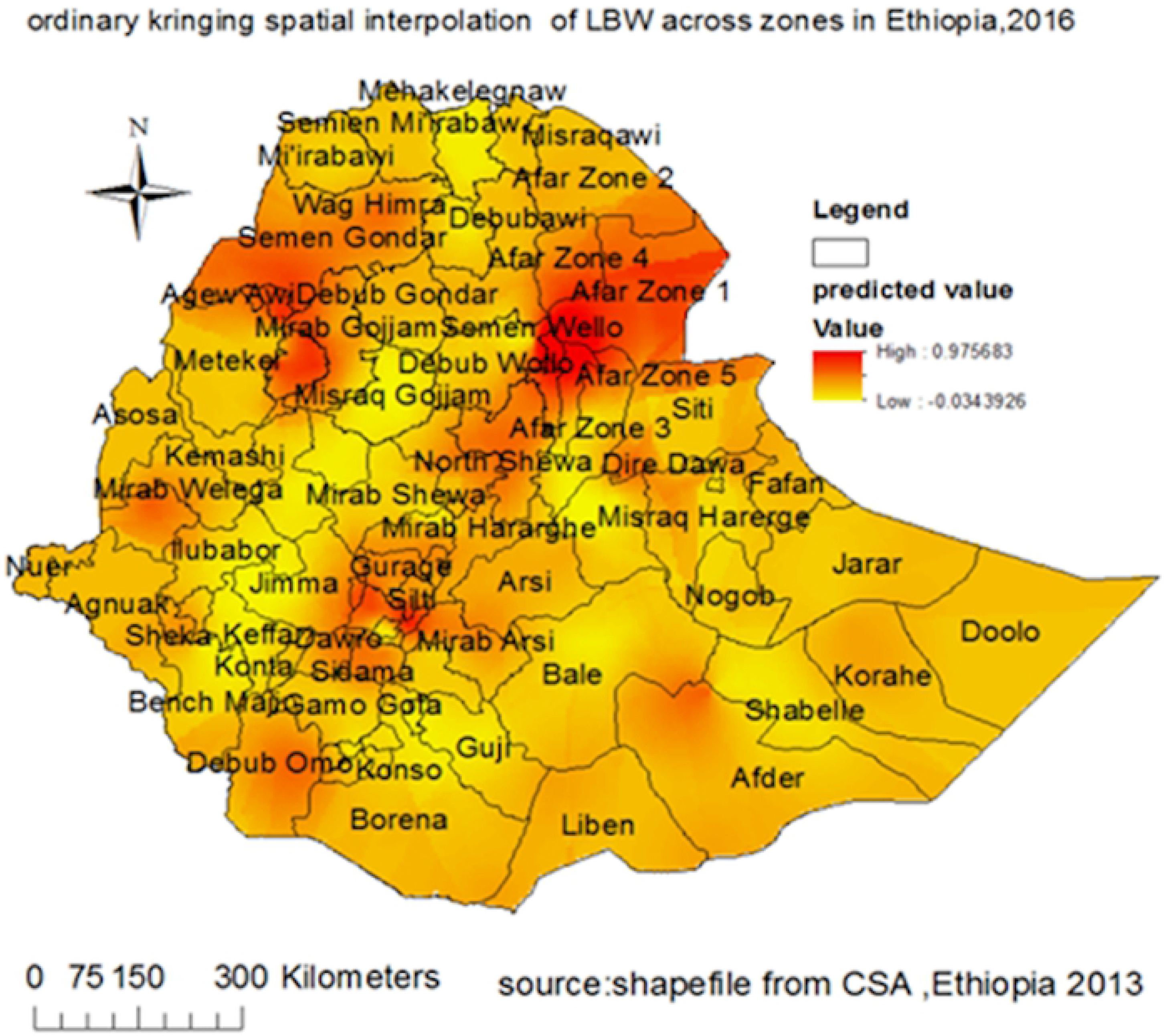
the ordinaiy kriging spatial interpolation of LBW across regions in Ethiopia, EDHS 2016. Continuous image produced by interpolating (ordinary Kriging interpolation method) LBW across different zones. TI1e dark red color indicates the predicted LBW high risk areas while dark greenish ramp color indicates less risk areas.

Fig 5: the ordinary kriging spatial interpolation of LBW across regions in Ethiopia, EDHS 2016.

### Multilevel mixed effects logistic regress analysis

#### Random effect analysis results

In the null model, variance component analysis was performed to decompose the total variance of LBW. The cluster-level variance which indicates the total variance of LBW that can be attributed to the context of the community in which the mothers were living was estimated. The applicability of multi-level mixed-effects logistic regression model in the analysis was justified by the significance of the community-level variance [community variance = 0.435; standard error (SE) = 0.19; P-value=0.001], indicating the existence of significant differences between communities regarding LBW incidence. The community variance was expressed as the intracluster correlation coefficient (ICC) and median odds ratio (MOR). The ICC was 0.117 which revealed that 11.7% of the total variance of LBW in Ethiopia can be attributed to the context of the communities where the mothers were dwelling. Since it was greater than 0.05 the nuisance of clustering was considered to produce reliable estimates (30). Moreover, the MOR was 1.88(95% CI 1.27, 2.34) which implied that the odds of having LBW was increased by 88% when mothers moved from low to high-risk communities.

In the full model community variance (community variance= 0.308; SE 0.17; P-value, 0.01), MOR 1.60 (95% CI 1.16, 2.14) and ICC (0.08) remained significant but reduced. About 8% of the total variance of LBW that can be attributed to the contextual-level factors remained significant even after considering some contextual risk factors for LBW. The PCV in this model was 29.2%. This showed that 29.2% of community variance observed in null model was explained by both community and individual level variables. Regarding model comparison, we used the loglikelihood ratio and deviance. The model with the highest loglikelihood or lowest deviance value (Model IV) was the best-fitted model. Cosequently, factors which are significant in this model were further considerd (Table 5).

**Table 4:** Random effects and model fitness

| Random effects | Model I | Model II | Model III | Model IV |
| --- | --- | --- | --- | --- |
| Community variance (SE) | 0.435*(0.187) | 0.323*(0.177) | 0.40*(0.18) | 0.308*(0.17) |
| ICC (%) | 11.70 | 9.0 | 10.0 | 8.0 |
| PCV (%) | reff | 25.3 | 8.0 | 29.2 |
| MOR (95%CI) | 1.88(1.27,2.34) | 1.72(1.10,2.16) | 1.83(1.23,2.28) | 1.70(1.16,2.14) |
| Model fitness | Model I | Model II | Model III | Model IV |
| Log likelihood | -732.80 | -701.00 | -729.20 | -688.09 |
| Deviance(-2LLR) | 1465.60 | 1402.00 | 1458.40 | 1376.18 |

\*significant at p-value <0.05;MOR:median odds ratio;ICC:intraclass correlation coefficient; PCV:proportional change in variance

**Table 5:** Multi level mixed effects logistic regression analysis of individual and community level factors associated with low birth weight in Ethiopia EDHS 2016.

| Characteristics | Model I | Model II | Model III | Model IV |
| --- | --- | --- | --- | --- |
| Fixed effects |  | AOR (95%CI) | AOR (95%CI) | AOR (95%CI) |
| Maternal age | - |  |  |  |
| 15-19 | - | 1.77(0.82,4.74) | - | 1.90(0.79,4.59) |
| 20-24 | - | 1.05(0.52,2.12) | - | 1.01(0.50,2.06) |
| 25-29 | - | 1.13(0.57,2.21) | - | 1.09(0.56,2.15) |
| 30-34 | - | 0.91(0.45,1.81) | - | 0.90(0.45,1.79) |
| 35-39 | - | 0.75(0.35,1.59) | - | 0.74(0.34,1.56) |
| 40-49 |  | 1 | - | 1 |
| Women educational level |  |  |  |  |
| No education | - | 1.91(1.23,2.93) | - | 1.82(1.12,2.96) * |
| Primary | - | 1.32(0.91,1.92) | - | 1.33(0.91,1.95) |
| Secondary and above | - | 1 | - | 1 |
| Gestational age |  |  |  |  |
| <37 weeks | - | 5.82(3.18,10.60) | - | 5.91(3.21,10.10) ** |
| ≥37 weeks | - | 1 | - | 1 |
| Iron |  |  |  |  |
| Yes | - | 1 | - | 1 |
| No | - | 1.15(0.84,1.57) | - | 1.14(0.83,1.57) |
| Number of ANC visit |  |  |  |  |
| <4 visits | - | 1.08(0.79,1.51) | - | 1.11(0.80,1.52) |
| ≥4 visits | - | 1 | - |  |
| Media exposure |  |  |  |  |
| Yes | - | 1 | - | 1 |
| No | - | 0.89(0.61,1.30) | - | 0.84(0.55,1.29) |
| Maternal BMI | - |  |  |  |
| Underweight | - | 0.94(0.62,1.42) | - | 0.96(0.63,1.45) |
| Normal |  | 1 |  | 1 |
| Overweight |  | 0.92(0.59,1.42) |  | 0.92(0.59,1.42) |
| Obese | - | 0.92(0.48,1.78) | - | 0.92(0.47,1.77) |
| Wealth index |  |  |  |  |
| Poor | - | 1.16(0.73,1.83) | - | 1.26(0.674,1.438) |
| Middle | - | 1.34(0.80,2.25) | - | 1.28(0.724,2.284) |
| Rich | - | 1 | - | 1 |
| Maternal anemia |  |  |  |  |
| Not anemic | - | 1 | - | 1 |
| Mild | - | 0.64(.33, 1.22) | - | 0.63(0.33,1.22) |
| Moderate | - | 1.07(0.56,2.05) | - | 1.09(0.57,2.08) |
| Severe | - | 1.48(1.04,2.11) | - | 1.47(1.04,2.01) |
| Sex of infant |  |  |  |  |
| Male | - | 1 | - | 1 |
| Female | - | 1.37(1.04,1.84) | - | 1.38(1.04,1.84)* |
| Community poverty level |  |  |  |  |
| Low | - | - | 1 | 1 |
| High | - | - | 0.93(0.59,1.47) | 0.79(0.47,1.35) |
| Community media exposure |  |  |  |  |
| Low | - | - | 1.34(0.81,2.02) | 1.20(0.76,1.89) |
| High | - | - | 1 | 1 |
| Community women education |  |  |  |  |
| Low | - | - | 0.79(0.52,1.20) | 0.83(0.52,1.30) |
| High | - | - | 1 | 1 |
| Place of residence |  |  |  |  |
| Urban | - | - | 1 | 1 |
| Rural | - | - | 1.09(0.71,1.68) | 1.01(0.62,1.60) |

AOR; adjusted odds ratio, CI; confidence interval, 1; referencecategory, \*: p-value<0.05, \*\*p-value<=0.01

#### Fixed effects analysis results

In the bi-variable mixed-effects logistic regression analysis wealth index, maternal age, number of ANC visit, gestational age, media exposure, mother’s education, maternal BMI, maternal anemia, sex of infant, iron supplementation during pregnancy, region place of residence, community poverty level, community illiteracy level, community media exposure and community women education were significant at p-value<0.2 and fitted for Multi-variable analysis.

Multivariable mixed effects logistic regression analysis was fitted to identify factors associated with low birth weight. In the final model (best fitted model to data) maternal education, maternal anemia, gestational age, and sex of the fetus were significantly associated with low birth weight. After keeping another individual, community level factors and random effect constant. The odds of bearing low birth weight baby among women who had no education was 1.82(AOR=1.82; 95% CI 1.12, 2.96) times higher as compared to women who had secondary and higher education. Regarding maternal anemia women who were severely anemic had 1.47 (AOR=1.47; 95% CI 1.04, 2.01) times higher odds of having low birth weight baby as compared to non-anemic mothers.

The likelihood of bearing low birth weight among women who had preterm delivery was 5.91(AOR=5.91;95%CI 3.21,10.10) times higher as compared to women who had term or post-term delivery. Women who had born female neonate had 38%(AOR=1.38;95%CI 1.04,1.84) increased odds of low birth weight baby delivery as compared to women who had born male neonate.

## Discussion

This study was conducted to assess spatial distribution and factors associated with low birth weight in Ethiopia. Thus, low birth weight was nonrandom in Ethiopia and affected by sociodemographic and pregnancy-related characteristics of mothers.

The spatial autocorrelation analysis revealed that low birth weight had spatial dependency (Moran’I=0.56, p-value: 0.01) in Ethiopia. This result was in agreement with studies done in upper- and middle-income countries(17, 31–33) where low birth weight had shown significant spatial clustering across different regions within respective countriess. This might be due to the variation in maternal health service utilization during pregnancy and variation in health service coverage across different regions in Etiopia.

The spatial scan statistics identified fifteen most likely clusters at the eastern part of Amhara and northern border of Benshangul gumuz region and 25 secondary clusters in the south Afar, southwest Amhara and northern part of Oromia region. The possible explanation could be the large disparity in health service aceess and affordability especially in those remote areas. This is supported by the fact that health service access is the major challenge in rural health in the countries where the majority of the population lives in rural areas like Ethiopia (34). Therefore,since SaTtscan is a powerful analytic tool (28) that identifies most likely clusters for intervention, especially in resource-limited areas, these spatial patterns and clustering of events, provide important information for the development and refinement of geographically based and population-specific prevention programs for maternal and child health to reduce LBW risk.

This study found that the odds of bearing LBW neonate higher among severely anemic mothers as compared to those without anemia. This finding is consistent with studies conducted in India (35, 36) and Ethiopia (37, 38). This might be due to the fact that anemia during pregnancy, especially if severe, could affect oxygen supply to the fetus and thus interferes with normal intrauterine growth or pregnancy duration which possibly leads to LBW (39).

The likelihood of LBW newborn delivery was higher among women without formal education as compared to women from secondary and above educational level. This finding was in agreement with studies conducted in Malawi (40), Bangladesh (22), India (21), and northwest Ethiopia (18). This may be due to the fact that uneducated mothers are relatively at low living standard and they might have poor maternal nutrition during pregnancy. In developing countries, it was found that poor gestational nutrition was found to be a major determinant of intrauterine growth restriction which might result in LBW delivery (39).

This study also identifies the odds of low birth weight among mothers who delivered before 37 weeks of gestation was higher as compared to those who delivered after 37 weeks of gestation. This result was concordant with studies conducted in Pakistan (41), Kenya (42), northwest (43), and southwest (19) Ethiopia. The possible explanation might be babies who were delivered in earlier periods of gestation were less likely to have full fetal development. Furthermore, evidence from the systematic review showed that gestational duration was found to be the most proximal cause of low birth weight (39).

The mothers who deliver female babies had higher odds of bearing low birth weight neonate as compared to those who deliver male neonates. This result is in line with the findings in Ghana (44) and Nepal (45). The association could be explained by the pathophysiologic mechanism in the uterus. Females had a higher risk of developing intrauterine growth restriction (IUGR) than males which probably results in low birth weight (39).

This study was based on the most recent EDHS data with a nationally representative large sample size based on a multilevel modeling approach. The sampling weight was applied to produce appropriate standard errors and then a reliable estimate. Despite the above strengths, the study had the following limitations. Some participants, data on birth weight was collected by mothers’ reports by recalling weight of their child at birth (recall bias) which may over or underestimate the results. Second, since it is secondary data analysis, those behavioral factors which will affect the out come of pregnancy were not included.

## Conclusion

Low birth weight was spatially clustered in Ethiopia. High-risk areas were identified in western and southeast parts of Amhara, the central and western part of Oromia, northeast BenshangulGumuz, southern part of Afar, northern part of South nation nationality and people’s regions. Regarding factors, being severely anemic, having no formal education, prematurity and bearing female neonate increase the odds of delivering low birth weight baby. Acordingly, focusing the policy intervention in those geogrsphically low birth weight risk areas and improving maternal education and nutrion (to reduce anemia) could be vital to reduce the low birth weight disparity in Ethiopia.

## Declarations

## Abbreviations

ANC: Antenatal care
AOR: adjusted odds ratio
EDHS: Ethiopian demographic and health survey
LBW: Low birth weight
MOR: median odds ratio
IUGR: Intrauterine growth restriction.

## Authors’ contributions

AML: Wrote the research proposal, conducted the data analysis, interpreted the results and organized the manuscript. AAM and MMS: Involved in designing the study, revising the proposal, guiding the statistical analysis, and write up of the manuscript. All authors read and approved the final manuscript.

## Acknowledgments

The authors would like to thank measure DHS for their permission to access the DHS datasets and central statistical agency for the shapefile.

## Availability of data and materials

The DHS data is publicly available and the authors approved the availability of data used for this study.

## Ethics approval and consent to participate

Ethical clearance was approved by an Institutional ethical Review committee of the Institute of Public Health, College of Medicine and Health Sciences and University of Gondar. Permission for data access was obtained from measure demographic and health survey through online request at http://www.dhsprogram.com. The authorization letter was also gained from measure DHS. Finnaly, No information obtained was disclosed to third body.

## Consent for publication

Not aplicable

